# Fused dorsal-ventral cerebral organoids model human cortical interneuron migration

**DOI:** 10.1101/131250

**Authors:** Joshua A Bagley, Daniel Reumann, Shan Bian, Juergen A Knoblich

**Affiliations:** Institute of Molecular Biotechnology of the Austrian Academy of Science (IMBA), Vienna 1030, Austria

## Abstract

Development of the forebrain involves the migration of GABAergic interneurons over long distances from ventral into dorsal regions. Although defects in interneuron migration are implicated in neuropsychiatric diseases such as Epilepsy, Autism, and Schizophrenia, model systems to study this process in humans are currently lacking. Here, we describe a method for analyzing human interneuron migration using 3D organoid culture. By fusing cerebral organoids specified toward dorsal and ventral forebrain, we generate a continuous dorsal-ventral axis. Using fluorescent reporters, we demonstrate robust directional GABAergic interneuron migration from ventral into dorsal forebrain. We describe methodology for time lapse imaging of human interneuron migration that is inhibited by the CXCR4 antagonist AMD3100. Our results demonstrate that cerebral organoid fusion cultures can model complex interactions between different brain regions. Combined with reprogramming technology, fusions offer a possibility to analyze complex neurodevelopmental defects using cells from neuropsychiatric disease patients, and to test potential therapeutic compounds.

## Introduction

The cerebral cortex contains two main populations of neurons, excitatory glutamatergic pyramidal neurons, and inhibitory γ-Aminobutyric acid (GABA) producing interneurons^1^. Excitatory cortical neurons are predominantly generated by dorsal forebrain progenitors, while inhibitory GABAergic cortical interneurons are generated by ventral forebrain progenitors^2^. To integrate into cortical circuits, interneurons perform a long-distance migration from their ventral origin into their target dorsal cortical regions^3^. This long-range tangential migration is controlled by many signaling pathways^4,5^, and studies using animal models indicate that mutations in some neurological disease-associated genes can disrupt interneuron migration^6,7^. However, the relationship between patient-specific mutations and human brain development remains enigmatic without suitable experimental human model systems.

Three-dimensional (3D) organoid culture technology allows the development of complex, organ-like tissues reminiscent of *in vivo* development^8^. Importantly, cerebral organoids recapitulate many aspects of embryonic cortical development including the generation of diverse cell types corresponding to different brain regional identities^9^. For instance, cerebral organoids can produce dorsal and ventral forebrain progenitors that generate excitatory neurons and inhibitory interneurons, respectively^9^. Moreover, cerebral organoids can be generated from human patient-derived induced pluripotent stem cells (hiPSCs), and used for functional genomic studies of neurological disorders such as microcephaly^9^ and Autism^10^. Therefore, cerebral organoids represent an exemplary experimental system to study the role of neurological disease-associated genes in brain development.

In the current study, we focused on expanding the cerebral organoid model system by enhancing the phenotypic analyses to include neuronal migration. We developed an organoid co-culture “fusion” paradigm to successfully combine independently patterned organoids into a single tissue. Using this approach, we recreated the dorsal-ventral forebrain axis, and by labeling one organoid with a fluorescent reporter, we observed robust migration of cells between fused organoids. The molecular taxonomy and migratory dynamics of these cells resembles that of cortical interneurons. Therefore, we developed an organoid fusion assay that allows analysis of human cortical interneuron migration. This technology enhances the repertoire of phenotypic assays available for cerebral organoids, and in turn the complexity of phenotypes that can be used to study the developmental cell biology of human neurological diseases.

## Results

### Drug-patterning of cerebral organoids enhances the production of ventral forebrain identity

The cerebral organoid method^9^ is capable of producing many different brain regions, including dorsal and ventral forebrain. As it relies on intrinsic patterning, however, the production of some regions is variable and infrequent, especially more ventral (NKX2-1+) interneuron progenitor regions (Renner at al, EMBO J, in press). To increase the consistency and yield of ventral-forebrain interneuron progenitor regions, we modified the organoid protocol^9,11^ to include a ventral drug-treatment (Figure 1A). Based on previous 2D-neuronal differentiation^12-14^ protocols, we utilized a combination of WNT-inhibition and enhanced SHH signaling to promote a rostro-ventral forebrain identity. qPCR analysis of the ventral organoids revealed a significant increase in expression of the forebrain marker FOXG1^15,16^ (Figure 1B,C). The dorsal forebrain marker TBR1^17^ became undetectable, while the ventral forebrain marker DLX2^18-20^ was dramatically increased in ventral organoids compared to control organoids (Figure 1B,C). To further confirm the successful ventralization of cerebral organoid tissue, we examined the expression of specific markers of ventral forebrain ganglionic eminence (GE) subregions that produce interneurons of different subtypes^3,21-23^. GSX2 is expressed in the dorsal-lateral GE (LGE) and caudal GE (CGE), NKX2-1 is expressed in the ventral-medial GE (MGE), while LHX6 is expressed in a subregion of the MGE producing more ventral-derived MGE (vMGE) interneurons ^3,20,21,24,25^ (Figure 1B). GSX2 was expressed in control organoids, but further increased in ventral organoids (Figure 1C). Expression of NKX2-1 and LHX6 was undetectable in control organoids, but largely increased in ventral organoids (Figure 1C). Finally, immunostaining confirmed the qPCR results indicating widespread expression of FOXG1 in control and ventral organoids, but only ventral organoids expressed the ventral forebrain marker NKX2-1 (Figure 1D). Intriguingly, control organoids widely expressed dorsal forebrain markers for both progenitors (PAX6)^26^ and early born neurons (TBR1)^17^ (Figure 1E). In contrast, ventral organoids contained only small regions of PAX6^+^ or TBR1^+^ tissue (Figure 1E). Therefore, these results confirm the successful production of ventral cerebral organoids at the expense of dorsal tissue upon ventral drug-treatment.

**Figure 1.**
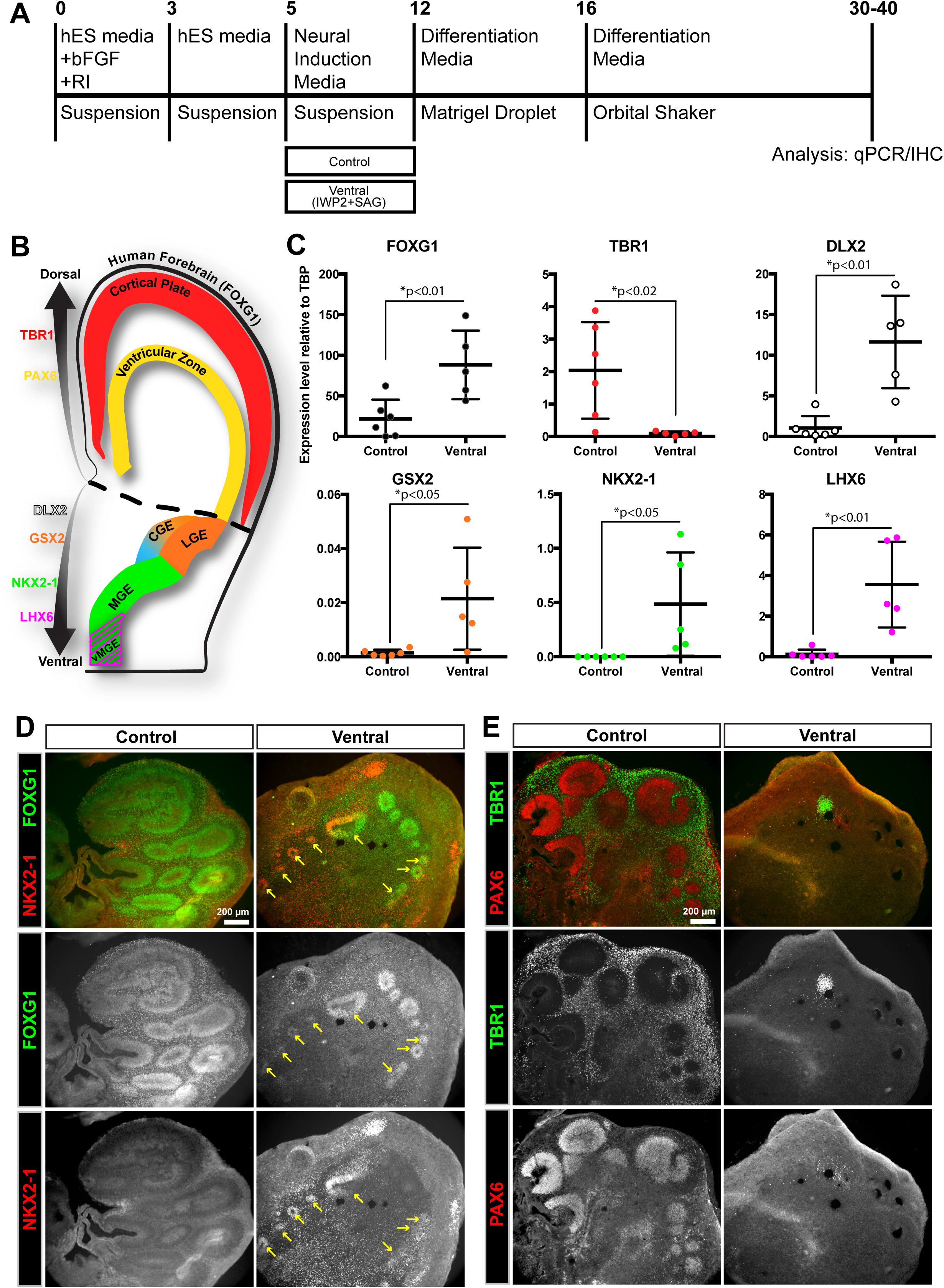
Ventral drug-treatment produces ventral-forebrain containing cerebral organoids. (A) Schematic of cerebral organoid protocol with ventral drug-patterning application (2.5μM IWP2 and 100nM SAG) during the neural induction step. (B) Schematic of a human coronal brain slice indicating the regional expression of the patterning markers used for qPCR/IHC analysis. (C) qPCR analysis of the expression of different brain regional markers showing an increase in ventral and decrease in dorsal forebrain identity in ventral cerebral organoids. Values are plotted as relative expression level (2^-ΔCt^) to the reference gene TBP. Each data point corresponds to a pooled batch of 8-10 organoids. Data is represented as mean±SD, and statistical significance was tested using the student’ s t-test (df=9) for control (n=6 batches) versus IWP2+SAG (n=5 batches) treatment. (D-E) Widefield images of immunostaining analysis of control and ventral cerebral organoids indicating the ventral-forebrain identity of ventral organoids (D), and the dorsal-forebrain identity of control organoids (E). Scale bars are 200μm.

### Fused cerebral organoids recapitulate a continuous dorsal-ventral forebrain axis and long-distance cell migration

To recreate the complete dorsal-ventral identity axis in a single tissue, we developed an organoid co-culture method, which we termed organoid “fusion”. Control organoids produced mostly dorsal forebrain tissue (Figure 1C,E). However, inhibition of SHH activity with the smoothened receptor inhibitor cyclopamine A (CycA) can enhance dorsal forebrain identity in 2D neuronal differentiation from hPSCs^27^. Therefore, to support dorsal identity, organoids were treated with CycA during the neural induction step of the cerebral organoid protocol (Figure 2A). In this approach, embryoid bodies (EBs) are individually patterned into either dorsal (CycA) or ventral (IWP2+SAG) forebrain organoids (Figure 2A,B). After patterning treatments, a ventral and a dorsal EB are embedded together within a single Matrigel droplet (Figure 2A), and over time the organoids grow together and become fused (Figure 2B). Immunostaining of fused ventral::dorsal organoids revealed the production of a continuous tissue where one side was highly positive for the ventral marker NKX2-1, while the opposite side was positive for the dorsal marker TBR1 (Figure 2C). Therefore, the organoid fusion method allows dorsal and ventral forebrain regions to be juxtaposed in an arrangement similar to that occurring during brain development.

**Figure 2.**
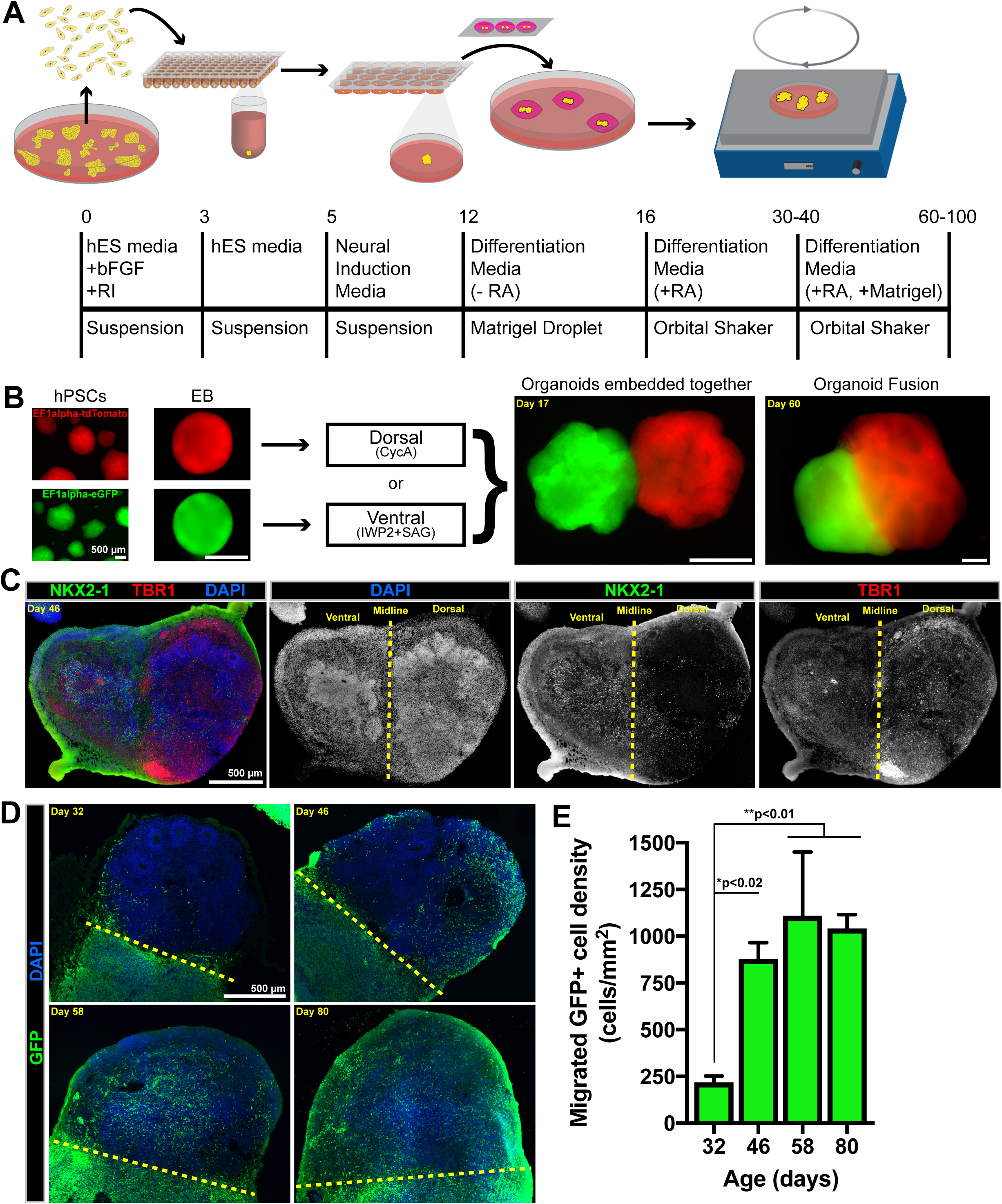
Fusion of cerebral organoids allows cell migration between ventral and dorsal forebrain tissue. (A) The experimental outline of the cerebral organoid fusion co-culture method. (B) Representative widefield images at different stages throughout the organoid fusion procedure. (C) Tile-scan image of an immunostained cryosection of a ventral::dorsal organoid fusion indicates the combination of ventral (NKX2-1^+^) and dorsal (TBR1^+^) regions. (D) Immunostained ventral::dorsal organoid fusion cryosections from organoids of different ages shows the time course of GFP^+^ cells migrating from ventral into dorsal tissue. (E) The GFP^+^ cell density in dorsal tissue was quantified in sections from 32 (n=3 organoids) 46 (n=3), 58 (n=4), and 80 (n=4) day-old organoids. The data is presented as mean±SD and statistical significance tested using the one-way ANOVA [F(3,10)=12.59, p=0.0010] with posthoc Tukey’ s test for between group comparisons. Scale bars are 500μm.

To test whether cells could migrate between the fused organoids we used cell lines containing a ubiquitous GFP reporter to create ventral/GFP^+^::dorsal organoid fusions. Immunofluorescent analysis showed many GFP^+^ cells from the ventral organoid within the GFP^-^ dorsal organoid (Figure 2D). Migrating cells were observed in small numbers around day 30 (Figure 2D,E). Their density drastically increased from day 30 to 46, but we did not observe a significant increase from day 46 to 80 (Figure 2E). Since the organoids increase in size with age, the absolute numbers of migrated cells must increase over time to maintain a similar density. Moreover, the cells appeared more dispersed throughout the dorsal regions in 80 day old organoids (Figure 2D). Therefore, based on these results, we focused our future analysis on organoids older than 60 days old.

### Mixing the tissue components of cerebral organoid fusions indicates directed ventral-to-dorsal cortical cell migration

During *in vivo* brain development, GABAergic interneurons originate in ventral forebrain progenitor regions before they migrate to their dorsal forebrain target^4,5^. To test whether this directionality is recapitulated in fused organoids, we varied the identities of their individual tissue components (Figure 3). We consistently labeled one organoid with GFP and the other with tdTomato, and delineated the fusions in an origin::target (GFP::tdTomato) arrangement. Using this paradigm, we analyzed the number of migrating cells when differentially controlling the dorsal/ventral identity of the origin and target regions of cerebral organoid fusions. Whole-mount imaging of ventral::control and ventral::dorsal fusions indicated the occurrence of GFP^+^ spots within the tdTomato^+^ tissue (Figure 3A). Conversely, the spots were rarely observed in control::control or dorsal::dorsal fusions (Figure 3A). This difference was even more striking when analyzing immunostained cyrosections of organoid fusion tissue. The highest amount of migrating cells occurred in ventral::control and ventral::dorsal fusions, while the least migration occurred in dorsal::dorsal fusions (Figure 3B,C). Although the average density of migrating cells in ventral::control fusions was not significantly different than ventral::dorsal fusions, the ventral::dorsal fusions more consistently contained a high amount of migrating cells (Figure 3C). These data confirm our initial observation of directionally biased migration from ventral into dorsal organoid tissue, and strongly suggests that the migration between fused organoids resembles interneuron migration. Finally, this experiment shows that ventral::dorsal fused organoids produce the most robust migration between organoids.

**Figure 3.**
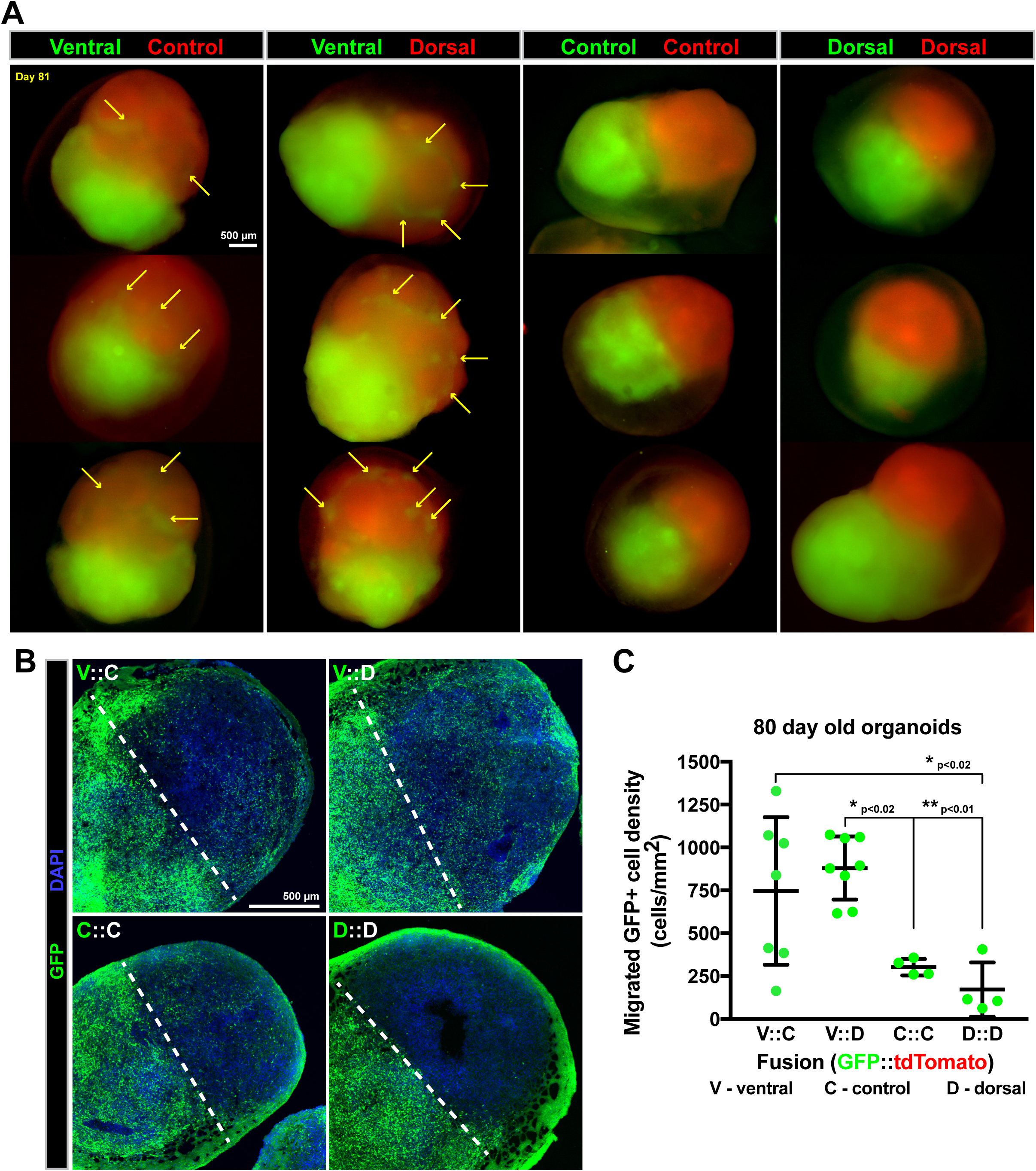
Mixing the tissue components of cerebral organoid fusions indicates the most robust migration from ventral into dorsal regions. (A) Cerebral organoid fusions were created containing different combinations of ventral (V), control (C) or dorsal (D) treated tissue. The components were labeled with either GFP (green) or tdTomato (red). (A) Whole mount images of ∼80 day old organoid fusions show the emergence of GFP^+^ spots (arrows) in tdTomato^+^ tissue in ventral-control and ventral-dorsal organoid fusions. (B) Tile-scan confocal images of immunostained mixed organoid fusion cyrosections shows migration of GFP^+^ cells across the midline (dashed line) into the GFP-organoid. (C) Quantification of GFP^+^ cell density in GFP^-^ tissue from tissue sections. Each data point corresponds to an individual organoid, and the data is represented as mean±SD with statistical significance tested using one-way ANOVA [F(3,19)=8.214, p=0.0010] with posthoc Tukey’ s test for between group comparisons. The ventral::control (n=7 organoids) and ventral::dorsal (n=8) fusions show the most migration of GFP^+^ cells compared to control::control (n=4) and dorsal::dorsal (n=4) fusions. Scale bars are 500μm.

### GABAergic interneurons migrate between fused cerebral organoids

Since our previous experiments suggested that the migration in ventral::dorsal organoid fusions resembles interneuron migration, we first tested whether the migrating cells were GABAergic by examining whether the migrating GFP^+^ cells expressed GAD1, one of the key enzymes for the synthesis of GABA^28^. Immunostaining revealed that the GFP^+^ cells that had migrated into the target organoid broadly expressed GAD1 (Figure 4A-C). Strikingly, when visualizing the entire organoid, GAD1 was expressed in a similar pattern as the GFP^+^ migrating cells (Figure 4A). Additionally, the expression of GAD1 appeared stronger in regions near the edge of the organoid (Figure 4B), and farther away from the origin of the migrating cells. Therefore, interneurons can migrate from ventral into dorsal regions within organoid fusions.

**Figure 4.**
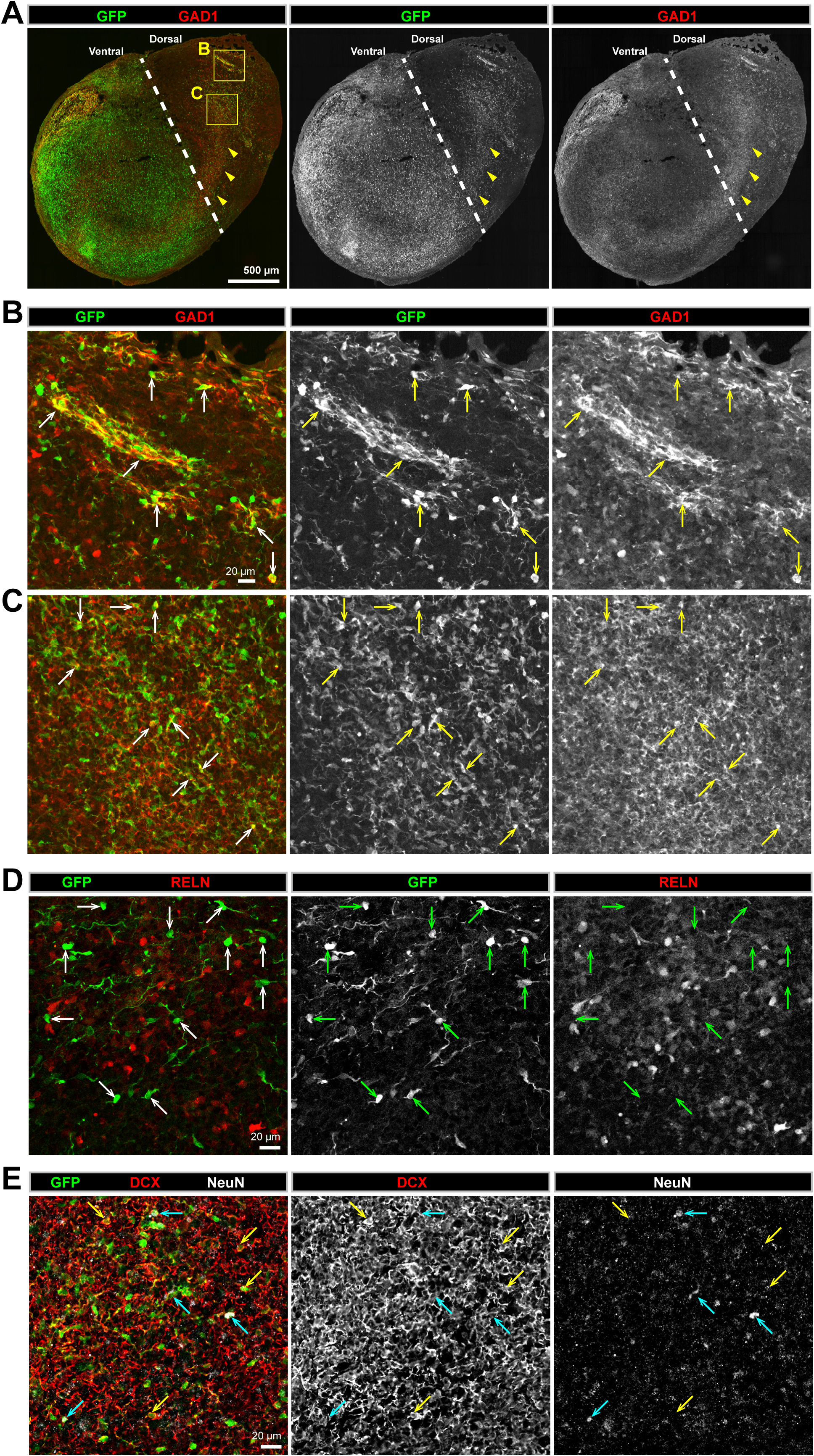
GABAergic interneurons migrate between fused dorsal-ventral cerebral organoids. (A) A whole organoid confocal tile-scan image of an immunostained 80-day old ventral::dorsal organoid fusion cryosection. GFP^+^ cells can be observed migrating across the fusion midline (dashed line) from GFP^+^ ventral into GFP^-^ dorsal tissue. The GABAergic marker GAD1 can be observed in a similar pattern as GFP (arrowheads). (B-C) A magnified view of peripheral (B) and internal regions (C) of the organoid fusion in (A). GFP^+^ cells expressing GAD1 can be observed in both regions (arrows). (D) A confocal image of GFP/RELN immunostaining in the dorsal region of an 80-day old ventral::dorsal organoid fusion cryosection showing that migrated GFP^+^ cells (arrows) do not express RELN. (E) A confocal image of GFP/DCX/NeuN immunostaining in the dorsal region of a 58-day old ventral::dorsal organoid fusion cryosection showing that migrating GFP^+^ cells are DCX^+^ immature neurons (yellow arrows), and some are mature (DCX^+^/NeuN^+^) neurons (blue arrows). Scale bars are (A) 500μm, (B-E) 20μm.

Another major population of tangentially migrating cells in the developing human brain are the non-GABAergic Cajal-Retzius cells^5^. These cells are identified by the expression of Reelin (RELN)^29,30^. Therefore, we tested whether the migratory GFP^+^ cells in organoid fusions also express RELN. Surprisingly, we observed an apparent lack of expression of RELN by GFP^+^ cells (Figure 4D), despite the substantial presence of GFP^-^, RELN^+^ cells in the dorsal target region of organoid fusions. Thus, our data indicate that the migratory GFP+ cells in organoid fusions are not Cajal-Retzius cells, and in combination with our previous results strengthens their identity as GABAergic interneurons.

Finally, we tested whether the migrating cells could produce mature neurons by analyzing their expression of the immature migrating neuron marker DCX^31^, and/or the mature neuronal marker NeuN^31^. We observed that the GFP^+^ cells expressed DCX (Figure 4E), confirming their neuronal identity, and a subset also expressed NeuN (Figure 4E). This finding indicates that the migrating GFP^+^ cells are migratory neurons (DCX^+^), but also that the migratory neurons can become mature neurons (DCX^+^/NeuN^+^). Taken together, our results show that interneurons can migrate between fused ventral-dorsal organoids, and can become mature neurons.

### Migrating interneurons produce various LGE/CGE and MGE-derived cortical interneuron subtypes in fused cerebral organoids

Since the migrating GFP^+^ cells can become mature neurons (Figure 4E), and appear to be predominantly GABAergic interneurons (Figure 4A,B), we next tested which interneuron subtypes are produced. Interneurons are particularly heterogeneous and multiple molecular markers can be used to identify various subtypes^32,33^ that are generated by distinct progenitor subpopulations within the ventral forebrain^3,20-23,34^. In humans, the majority of interneurons are generated from NKX2-1^+^ regions of the MGE^35,36^. Therefore, we tested whether the migrating GFP^+^ cells produce MGE-derived interneuron subtypes. In humans, SOX6 is expressed in the MGE and in immature and mature interneurons emerging from this region^35^. In organoid fusions, we observed multiple SOX6^+^ MGE interneurons among the GFP^+^ migrating cells GFP^+^ cells that were also GAD1^+^ (Figure 5A). Therefore, MGE-derived interneurons can be generated within cerebral organoid fusions. To confirm this finding, we also examined the expression of markers for MGE-derived interneurons^3,33,34^. We observed expression of somatostatin (SOM) (Figure 5B), neuropeptide Y (NPY) (Figure 5C) calbindin D-28k (CB) (Figure 5D), and parvalbumin (PV) (Figure 5E) by GFP+ migrated interneurons (GAD1+ or VGAT+). Therefore, in organoid fusions, multiple MGE-derived interneuron subtypes can be generated.

**Figure 5.**
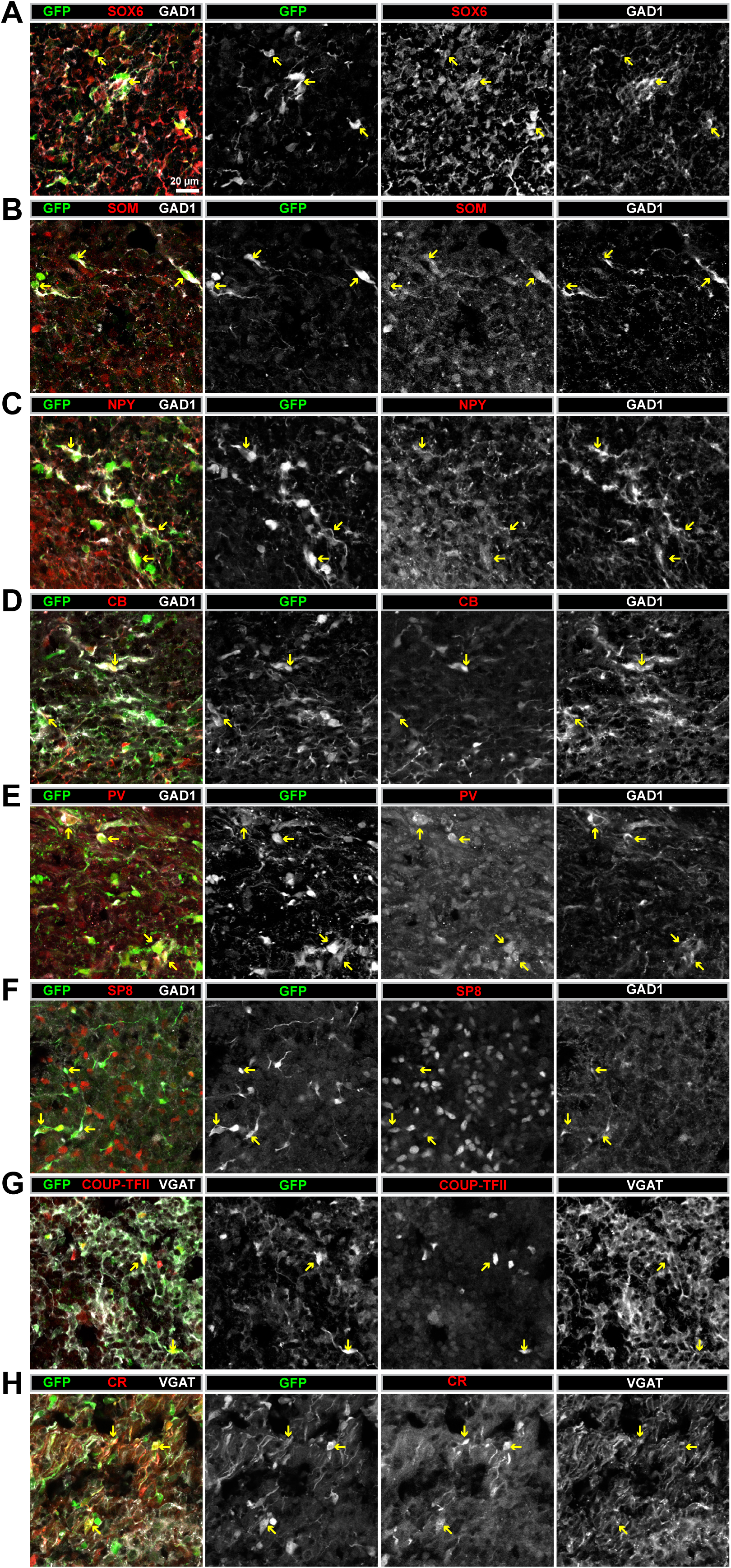
Migrating interneurons in ventral::dorsal cerebral organoid fusions express various interneuron subtype markers. (A-H) Confocal images of immunostaining in dorsal regions of 80-day old ventral::dorsal organoid fusion cryosections. Expression of the GABAergic markers GAD1 or VGAT were used to identify interneurons. Examples of various migrated GFP^+^ interneurons expressing either GAD1 or VGAT were observed expressing the MGE-derived interneuron marker SOX6 (A) or subtype markers SOM (B), NPY (C), CB (D), and PV (E). Migrated GFP^+^ interneurons also expressed the LGE/CGE-derived interneuron markers SP8 (F), COUP-TFII (G), or the subtype marker CR (H). Scale bars are 20μm. Abbreviations: SOM=somatostatin, NPY=neuropeptide Y, CB=calbindin, PV=parvalbumin, CR=calretinin, VGAT=vesicular GABA transporter, GAD1=glutamate decarboxylase 1.

The remaining interneurons in the human brain arise from the LGE/CGE^3,33,34^. Similar to SOX6 for MGE, the transcription factors COUP-TFII/NR2F2 and SP8 can be used as LGE/CGE fate-mapping markers for cortical interneurons^35^. Both SP8 (Figure 5F) and COUP-TFII (Figure 5G) were expressed by GFP^+^ migrating interneurons (GAD1^+^) in organoid fusions. As confirmation of this result, we also analyzed the expression of markers for LGE/CGE-derived subtypes. Our previous data already indicated a lack of RELN expression by GFP^+^ migrating cells (Figure 4), which in addition to non-interneuron Cajal-Retzius cells is also expressed by subpopulations of GABAergic MGE and CGE-derived interneurons^3,34^. We also did not observe any vasoactive intestinal peptide (VIP) expressing GFP^+^ interneurons (data not shown). However, we did observe calretinin (CR) expressing GFP^+^ interneurons (VGAT^+^) in organoid fusions (Figure 5H). Collectively, these results indicate that organoid fusions contain many diverse interneuron subtypes originating from the major ventral forebrain subregions (MGE and LGE/CGE).

### Neuronal migration in fused cerebral organoids resembles the tangential migration of cortical interneurons

The migratory dynamics of tangentially migrating cortical interneurons has been extensively documented using time-lapse imaging experiments^37-42^. This behavior differs from that of radially migrating cortical cells which move along a radial glia fiber, or that of linearly migrating cells, such as the chain migration observed in LGE-derived olfactory bulb interneurons^5,43^. Therefore, to determine whether the behavior of migrating GFP^+^ cells in organoid fusions resembles that of cortical interneurons, we performed time-lapse recordings of migrating GFP^+^ cells in ventral-dorsal organoid fusion slice-cultures (Figure 6A). The GFP^+^ ventral origin region of the organoid fusion could easily be distinguished from the GFP^-^ dorsal target region (Figure 6A). Thus, we could easily visualize the morphology of the sparsely labeled GFP^+^ cells which had migrated into the dorsal organoid fusion region. Over 3 days, we observed both stationary and motile cells (Figure 6B, Supplemental movies 1-5), and surprisingly we also observed dynamic neuronal processes resembling axons that extended long distances (Figure 6C, Supplemental movie 6). Strikingly, the migratory cells exhibited dynamics (Supplemental Movies 1-6) which exactly resembled the dynamics of migrating interneurons in the marginal zone (MZ) and intermediate zone (IZ) of embryonic mouse cortex^37-42^. Migrating cells contained leading and trailing processes. The leading process was usually branched, and the migratory direction was determined by the branch dynamics. For instance, we observed multiple examples of cells extending a branch in the future direction of travel while retracting a branch that was extended in an alternate direction (Figure 6B). In addition, cells migrated in multiple directions with frequent abrupt changes in direction, which resembles the multidirectional, wandering behavior exhibited by tangentially migrating cortical interneurons within the cortex ^42^.

**Figure 6.**
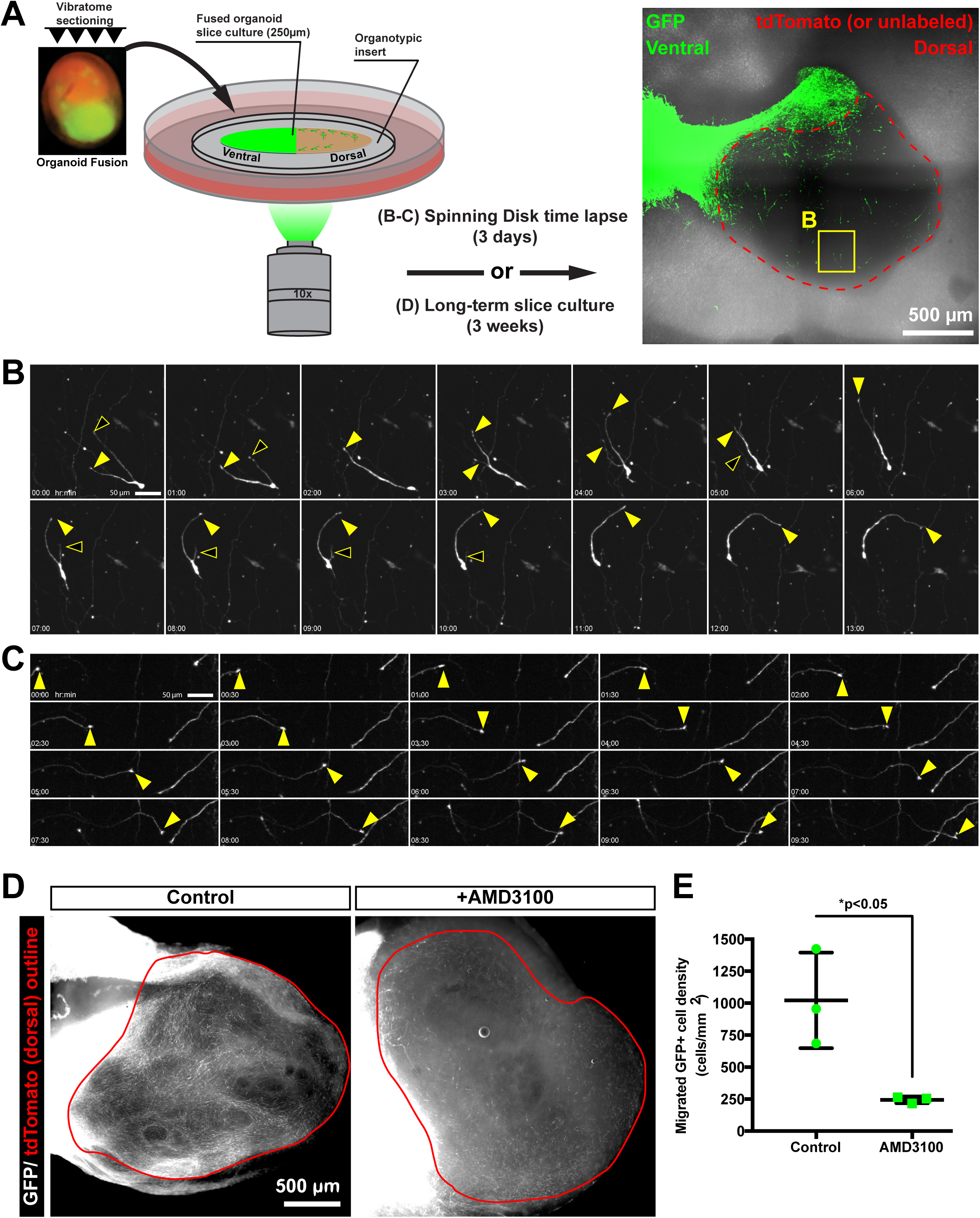
Migrating cells in cerebral organoid fusions exhibit the migratory dynamics of tangentially migrating interneurons and are sensitive to CXCR4 activity. (A) A schematic representation of the organoid fusion slice culture assay for either short-term time-lapse imaging of migratory dynamics, or long-term drug-treatments. A representative tile-scan image of an entire ventral::dorsal organoid fusion slice is shown with the ventral GFP^+^ regions labeled in green and the unlabeled or tdTomato dorsal regions outlined in red. A region containing the migrating cell shown in (B) is noted by a yellow box. (B) Still images from a 3-day time-lapse experiment showing a migrating GFP^+^ cell. The branched leading process exhibits both extending (closed arrowheads) and retracting (open arrowheads) branches as the cell body follows one of the leading processes. (C) An extending neurite (closed arrowhead) with a tuft that appears to be an axon growth cone travels in one direction across the field of view. (D) A widefield image of GFP^+^ cells that migrated into the tdTomato^+^ dorsal region (red outline) from long-term organoid fusion slice cultures that were either untreated (control) or treated with a the CXCR4 inhibitor (AMD3100). (E) Quantification of the migrated GFP^+^ cell density with data represented as mean±SD with statistical significance tested using the student’ s t-test (df=4) comparing control (n=3 organoids) to AMD3100 (n=3) treatment. Each data point represents a one slice from individual organoids. Fewer migrating cells are observed in AMD3100-treated slice cultures. Scale bars are (A) 500μm, (B-C) 50μm, and (D) 500μm.

The chemoattractant SDF-1 (CXCL12) and its receptor CXCR4 regulate the tangential migration of interneurons^44^. To test if the migration we observed in cerebral organoid fusions is CXCR4-dependent, we created long-term slice cultures of organoid fusions that we treated with the CXCR4 antagonist, AMD3100. Compared to untreated control slices prepared from the same organoid fusion, the density of migrating GFP^+^ cells into GFP^-^ dorsal regions was significantly reduced upon AMD3100 treatment. Therefore, the cell migration in cerebral organoid fusions depends on CXCR4 activity. Combined with our previous data, this result confirms that the cell migration observed in cerebral organoid is consistent with that of tangentially migrating interneurons.

## Discussion

In this study, we developed a cell migration assay using fused cerebral organoids. The concept of fusing organoids is based on classical co-culture experiments. However, our method represents an adaptation and extension of this technique to study human neuronal cell migration with the experimental control allowed by *in vitro* 3D cell culture. Ventral-dorsal organoid fusions exhibit robust long-distance cell migration resembling that of ventral forebrain-derived cortical inhibitory interneurons. This is based on multiple pieces of data. First, we observed substantially more migration from ventral into dorsal forebrain regions in cerebral organoid fusions. Second, the migrating cells within organoid fusions express GABAergic markers (GAD1/VGAT). Third, the migrated GFP^+^ cells can produce a variety of interneuron subtypes. In addition, the lack of RELN expression by migrating GFP^+^ cells supports a ventral forebrain-derived interneuron identity. Fourth, the migration dynamics of the GFP^+^ cells in fused organoids resembles the characteristic migratory behavior exhibited by interneurons migrating tangentially within the cortex. Finally, the cell migration in organoid fusions was inhibited by CXCR4 inhibition as observed for tangentially migrating interneurons in developing mouse cortex. Therefore, we have recapitulated the migration of human cortical interneurons from ventral in dorsal forebrain regions. Moreover, the observance that these cells exhibit both immature and mature neuronal markers indicates their ability to undergo maturation once arriving in their dorsal cortical forebrain target regions. This represents the exciting opportunity to recreate a developing human cortical circuit containing an enhanced repertoire of cellular diversity than was possible to achieve within the previous singular organoid method.

An exciting future application of this technique will be to study human developmental biology and its relationship to neurological diseases. With this in mind while characterizing this assay, we present different experimental paradigms, each of which can be used to investigate various aspects and scientific questions related to neuronal cell migration. For instance, the labeling of the cells within one of the fused organoids with a fluorescent reporter allows the visualization of long-distance neuronal migration. This can even be continuously monitored over time through whole-mount imaging of intact organoid fusions. In addition, the identity of the cells can be easily examined using immunostaining for identification of neuronal subtypes. This is useful because many psychiatric diseases, such as Schizophrenia, are thought to involve selective deficits in specific interneuron subpopulations^45^.

A second type of analysis is dissecting the role of specific molecules on neuronal migration to determine if they are acting in a cell-autonomous or cell non-autonomous manner. For instance, using our organoid mixing paradigm, organoids from either the origin of migration or the target can be genetically manipulated independently by deriving the origin and target organoids from distinct cell lines containing either mutant or wild-type alleles of a gene of interest. This is a powerful genetic tool routinely used in genetic model organisms, but can now also be utilized in 3D cultures of developing human brain tissue.

We also present an organoid fusion slice culture paradigm in which confocal time-lapse imaging can be used to analyze the short-term dynamics of neuronal cell migration. This is an important tool because some molecules such as GABA are known to affect the motility of migrating interneurons^4^, which may produce a subtle phenotype in a long-term migration assay, and therefore instead require a higher time-resolution analysis of the dynamics of migrating cells. In addition, long-term slice cultures can be used to test how different drug-treatments can affect cell migration. This presents the opportunity to perform drug-screens to determine the effect of many different molecules on interneuron migration.

Finally, the organoid fusion method also presents unique applications beyond cell migration, which can be developed further in the future. As one example, we observed substantial neurite dynamics with the appearance of axon projections. In addition, although we specifically focused on ventral-dorsal forebrain fusions, the organoid fusion paradigm allows flexibility in the brain regional identities that can be grown together. Combined with the additional brain-region specific organoid protocols^46-49^, the possible brain circuits that could be modeled using organoid fusions is vast. Therefore, organoid fusion technology greatly enhances the phenotypic analyses possible in cerebral organoids, and the flexibility of the method immensely expands the future development of *in vitro* models of human neurological diseases.

## Methods

### Cell culture

Feeder-dependent human induced pluripotent stem cells (hiPSCs) (Systems Biosciences, cat. no. SC101A-1) were obtained from System Biosciences with pluripotent verification and contamination-free. Feeder-dependent hiPSCs were cultured with irradiated mouse embryonic fibroblast (MEF) feeder cells (MTI-GlobalStem, cat. no. 6001G) on gelatin coated (0.1% gelatin in PBS) 6-well culture plates using human embryonic stem cell (hESC) medium: DMEM/F12 (Invitrogen) containing 20 ng/mL bFGF (produced by IMBA institute Molecular Biology Service core facility), 3.5μL/500mL media of 2-mercaptoethanol, 20% KnockOut Serum (Invitrogen), 1% GlutaMAX (Invitrogen), 1% MEM-NEAA (Sigma), and 3% FBS). Feeder free H9 human embryonic stem cells (hESCs) were obtained from WiCell with verified normal karyotype and contamination-free. Feeder-free hESCs were cultured on hESC-qualified Matrigel (Corning cat. no. 354277) coated 6-well plates using mTeSR1 (Stemcell Technologies). All stem cells were maintained in a 5% CO_2_ incubator at 37°C. Standard procedures were used for culturing and splitting hPSCs as explained previously^11^. All cell lines were routinely tested for contamination and confirmed mycoplasma-negative.

### Cloning/Molecular biology/Generating hPSC lines

For ubiquitous fluorescent labeling of cells, a reporter construct was inserted into the safe-harbor AAVS1 locus in hPSCs as done previously with TALEN technology ^50^ using the AAVS1 SA-2A-Puro donor vector (gift from Dr. Rudolf Jaenisch) as a template. A modified backbone was created containing flanking tandem repeats of the core chicken HS4 insulator (2xCHS4). Fluorescent reporter expression cassettes were inserted between the flanking insulator sequences. Tthe following expression constructs were inserted into iPSCs: 1) 2xCHS4-EF1α-eGFP-SV40-2xCHS4, 2) 2xCHS4-EF1α-tdTomato-SV40-2xCHS4. In feeder-free H9 hESCS, a 2xCHS4-CAG-eGFP-WPRE-SV40-2xCHS4 construct was inserted to enhance the GFP expression for time-lapse imaging experiments.

hPSCs were prepared for nucleofection as a single-cell suspension using the same cell dissociation procedures as for making EBs^11^. The Amaxa nucleofector (Lonza) was used with Stem Cell Kit 1 using 800,000 cells per nucleofection containing 1μg each of the TALEN guides, and 3μg of the donor plasmid DNA following manufacturer guidelines. After nucleofection, 200μL of the total 600 μL nucleofection solution was plated onto a 10 cm cell culture dish. Colonies from single cells were grown for 4 days, and then treated with puromycin (Puro) (Jena Bioscience, cat. no. NU-931-5). For feeder-dependent cells, 1.1 μg/mL puro was applied, and for feeder-free cells 0.7 μg/mL was applied. The puro treatment continued for 5-7 days until the surviving colonies were large enough to be picked manually and transferred into a 24-well plate. When splitting the colonies from a 24-well plate, half of the cells were used for genotyping, while the other half was expanded into 12 and then 6-well formats, and then used for further experiments. For genotyping, DNA was extracted using the QuickExtract solution (EpiCentre), and PCR was performed using the following primers to identify correctly targeted AAVS1 insertions: 1) Puro (*AAVS1_Puro-fwd*: tcccctcttccgatgttgag and *AAVS1_Puro-rev*: gttcttgcagctcggtgac), 2) eGFP (*AAVS1_eGFP-fwd*: GAACGGCATCAAGGTGAACT and *AAVS1_eGFP-rev*: cttcttggccacgtaacctg), and 3) tdTomato (*AAVS1_tdTomato-fwd*: ctacaaggtgaagatgcgcg) and (*AAVS1_tdTomato-rev*: tccagcccctcctactctag). To determine whether the insertion was heterozygous or homozygous, the presence of a WT allele was tested using additional PCR primers: 4) WT (*AAVS1_WT-fwd*: tcccctcttccgatgttgag, and *AAVS1_WT-rev*: tccagcccctcctactctag). Cell clones with correctly targeted heteroyzygous or homozygous insertions were archived by freezing with Cell Banker 2 solution (Amsbio, cat. no. 11891) and/or cultured for further experiments.

### Cerebral organoid generation and fusion

Cerebral organoids were generated following the previous protocol developed in our lab ^9,11^ with slight modifications. A drug-patterning treatment was applied during the neural induction step of the protocol (∼day 5-11) using one of the following treatments: 1) Control, no drugs, 2) Ventral, 2.5μM IWP2 (Sigma, cat. no. I0536) and 100nM SAG (Millipore, cat. no. 566660), 3) Dorsal, 5μM CycA (Calbiochem, cat. no. 239803). Stock drug solutions were created as follows: IWP2 (5mM in DMSO), SAG (1 mM in H_2_O), and CycA (5 mM in DMSO). Following embedding in Matrigel (Corning, cat. no. 356235), organoids were grown in 10 cm cell culture dishes containing 25 mL of differentiation medium, and after the first media exchange maintained on an orbital shaker with medium exchange every 5-7 days. After day 40 of the protocol, when organoids begin to grow out of the Matrigel droplet, the differentiation medium was supplemented with 1% Matrigel.

To create organoid fusions, EBs were grown separately and individually patterned using either control, dorsal, or ventral protocols as described above. During Matrigel embedding, two EBs were transferred into the same parafilm well and embedded in a single droplet (∼30 μL) of Matrigel. The EBs were gently pushed as close together as possible using a 20 μL pipet tip to ensure the EBs would remain in close proximity within the middle of the solidified Matrigel droplet.

### RNA extraction/qPCR

For each drug-patterning treatment group, 8-12 organoids were collected at day 30-40 into 2 mL RNAse-free tubes and chilled on ice throughout the procedure. The organoids were washed 3x in cold PBS, then the Matrigel was dissolved by incubating the organoids in chilled Cell Recovery Solution (Corning, cat. no. 354253) for 1 hr at 4°C. The dissolved Matrigel was removed by rinsing 3x in cold PBS. RNA was extracted using the RNeasy mini kit (Qiagen). cDNA synthesis was performed using 2 μg of total RNA and the Superscript II (Invitrogen) enzyme according to the manufacturer protocols. qPCR reactions were performed using Sybr Green master mix (Promega) on a BioRad 384-well machine (CXF384) using the following reaction protocol: 1) 95°C for 3min, 2) 95 °C for 10s, 3) 62 °C for 10s, 4) 72 °C for 40s, 5) go to 2, 40 cycles, 6) 95°C for 1min, 7) 50 °C for 10s. Quantification was performed in excel by calculating the ΔCt value using TBP as a reference gene. Data is presented as expression level (2^-ΔCt^) relative to TBP.

### Cerebral Organoid Slice Culture and Drug-treatment

Slice cultures were generated by vibratome sectioning organoid fusions. The organoid fusion tissue was embedded in 4% low melt agarose (Biozym, cat. no. 850080), and sectioned in ice-cold PBS (without Ca^2+^/Mg^2+^) to create 200-250 μm sections. The sections were transferred onto Millicell organotypic inserts (Millipore, cat. no. PICM01250) in 6-well cell culture plates. For time-lapse imaging, sections were cultured for 1-2 days before mounting for imaging using a spinning disk confocal microscope (VisiScope). Sections were cut away from the culture insert membrane and inverted onto a glass-bottom dish. The section was immobilized by placing a cell culture insert on top of the section, and attaching it to the dish using vacuum grease. For drug-treatment, long-term slice cultures were initially cultured overnight in differentiation media (+5% FBS). Then the media was exchanged for fresh media (control) or media containing the CXCR4 inhibitor AMD3100 (Sigma, #A5602) and cultured for an additional 3 weeks before tissue fixation and further immunofluorescent processing.

### Histology/Cryosectioning/Immunofluorescence

Organoid tissue was collected at the desired age, rinsed 3x in PBS, and fixed in 4% PFA overnight at 4°C. The next day the tissue was rinsed 3x in PBS, and cryoprotected in 30% sucrose in PBS overnight at 4°C. Then the tissue was incubated 2-4 hours in a 1:1 mixture of 30% sucrose/PBS and O.C.T. cryoembedding medium (Sakura, cat. no. 4583). Next, groups of 2-4 organoids were transferred from the sucrose/OCT mixture into a cryomold and filled with O.C.T. The embedded tissue was frozen on dry ice, then placed in −80°C for long term storage until cryostat sectioning. Frozen organoid tissue was sliced into 20 μm sections using a cryostat (Leica), and collected on superfrost Ultra Plus slides. Tissue sections were arranged such that every 10^th^ slice was collected sequentially until each slide contained 8-10 sections per block of tissue. The sections were dried overnight, and then used for immunofluorescent labeling, or stored at −20°C until ready to stain.

Immunofluorescence was performed on tissue sections directly on slides. The O.C.T. was removed by washing 10 minutes in PBS using a slide rack (Sigma, Wash-N-Dry). The sections were post-fixed directly on the slides using 4% PFA for 10 minutes at RT followed by washing 3×10 minutes in PBS. The tissue area was outlined using a hydrophobic PAP pen. Permeabilization/blocking was performed using 5% BSA/0.3% TX100 in PBS with 0.05% sodium azide and incubated 30 minutes at room temperature (RT) within a dark humidified slide box. Primary antibodies (a table of used primary and secondary antibodies can be found at the end of the method section) were added at the desired dilutions in antibody solution (5% BSA, 0.1% TX100 in PBS with 0.05% sodium azide) and incubated overnight at 4°C. Slides were rinsed 3x in PBS and then washed 3x10 minutes in PBST at RT on an orbital shaker. Secondary antibodies were added at 1:500 in antibody solution and incubated at RT for 2 hours. DAPI solution (2 μg/mL) was added for 5 minutes, then slides were washed as done after primary antibody application, with the final washing using PBS. Coverslips were mounted using DAKO mounting medium (Agilent Pathology Solutions, cat. no. S3023) and allowed to harden overnight. Slides were then stored at 4°C until imaging, or at −20°C for long-term storage.

For long-term slice cultures, slices grown on cell culture inserts were rinsed in PBS, and then fixed in 4% PFA for 2 days at 4°C. The sections were washed 3x10 minutes in PBS, then permeabilized/blocked as performed for cryosections on slides. Primary and secondary antibody incubations were performed overnight at 4°C, followed by washing 3x10 minutes in PBS after each step. The sections were mounted in Vectashield containing DAPI (Vector Labs).

### Imaging/Microscopy

Fluorescent cell culture imaging was performed using a Zeiss Axio Vert.A1 widefield microscope and an Axiocam ERc 5s camera (Zeiss, Zeiss GmbH) using Zeiss Plan-Neofluar 2.5x 0.085 and Zeiss LD A-Plan 10x 0.25 Ph1 objectives. Both tdtomato and GFP channels were recorded separately and subsequently pseudocolored and merged using the Fiji package of ImageJ.

Widefield imaging of IHC stainings and long-term slice cultures was performed using an Axio Imager Z2 (Zeiss, Zeiss GmbH) using a Sola SM2 illumination source, (5x 0.15 plan-neofluar, 10x 0.3 plan-neofluar, 20x 0.5 plan-neofluar) objective and a Hamamatsu Orca Flash 4 camera. Filters used were Ex360/40nm Em 445/50nm, Ex480/40nm Em 535/50nm and Ex560/55nm Em 645/75nm.

Confocal imaging was performed using a Zeiss LSM700 AxioImager with a 20x 0.8 plan-apochromat dry objective. Lasers of 405nm (5mW), 488nm (10mW), 555nm (10mW) and 639nm (5mW) together with, corresponding to wavelength, filters SP490, SP555, SP640 and LP490, LP560, LP640 were used for recording. For whole organoid tile scans the XY Scanning Stage and the Zeiss Zen implemented stitching algorithm was used. For colocalization of markers, Z-scans at 20x were performed.

Imaging of GFP stained slides used for cell density counting, and time-lapse migration were acquired using a Yokogawa W1 spinning disk confocal microscope (VisiScope, Visitron Systems GmbH, Puchheim, Germany) controlled with the Visitron VisiView software and mounted on the Eclipse Ti-E microscope (Nikon, Nikon Instruments BV) using ex 488 laser and em filter 525/50. Measurements were recorded using the 10x 0.45 CFI plan Apo lambda (Nikon, Nikon Instruments BV) objective with the sCMOS camera PCO edge 4.2m. Whole IHC slice imaging for cell counting was performed using the tile scan acquisition function. Tile scans were then stitched using the Grid/Collection stitching plugin 51 in Fiji. For time lapse movies, tile scan Z-stacks were recorded. After stitching as performed for cell counting, a maximum Z-projection was created, and the Z-projection time stacks used for global visualization of cell migration. Cropped regions were created using Fiji and saved as uncompressed AVI files. The AVI files were converted to the mp4 format using Handbrake software to generate the supplemental movies.

For generation of figures and presenting images, the “crop” function, and changing the max/min levels using the “Brightness/Contrast” function of Fiji were used.

### Cell density quantification

The number of GFP^+^ cells migrating into GFP^-^ target regions of cerebral organoid fusions was counted manually using the “Cell Counter” plugin of Fiji. The GFP^-^ area in organoid fusions was outlined with the ROI tool and the area calculated using the “measure” function. The cell counts were divided by the area to determine the density of migrated GFP^+^ cells. For the migration time course (Figure 2E,F) and organoid fusion mixing (Figure 3B,C) experiments, confocal tile scan images of tissue cryosections were used for quantification. For the long-term slice culture experiment (Figure 6D,E), a wide-field 5x image containing the GFP^-^ region was used for counting.

### Statistics

All graphs and statistical analyses were generated using Prism 7 (GraphPad). When comparing only two groups (Figure 1C, 6E) an unpaired two-tailed Student’ s t-test was performed, for the remaining data comparing multiple groups a one-way ANOVA with posthoc Tukey’ s test was used (Figure 2F, 3C). Samples were tested for normality prior to testing statistical significance. No statistical methods were used to predetermine the sample size. Sample sizes for experiments were estimated based on previous experience with a similar setup that showed significance. Experiments were not randomized and investigator was not blinded.

## Data Availability

The authors declare that all data supporting the findings of this study are available within the paper, and its supplementary information files.

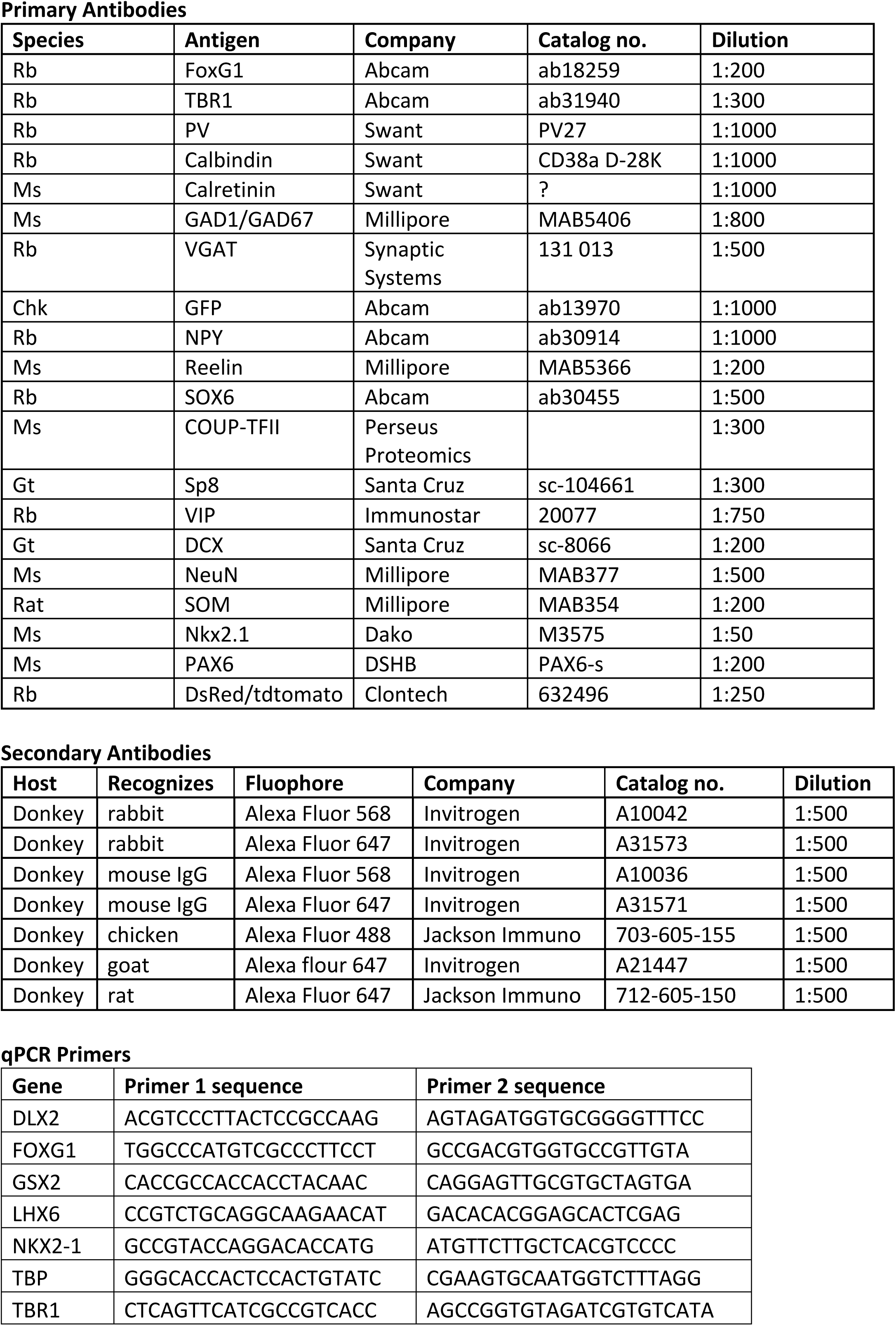

## Figure Legends

**Supplemental Movie 1:** A time-lapse movie of migrating GFP^+^ cells within the dorsal region of a ventral/GFP::dorsal organoid fusion. The cell migrates in a single direction. The leading process is branched with the different branches dynamically extending and retracting seemingly independent of one another. The trailing process follows as the cell body moves forward, and multiple times a leading process becomes a trailing process. As the cell moves forward, one leading process is extended while the remaining processes retract. Then the whole migratory dynamic cycle is repeated as the cell progresses forward.

**Supplemental Movie 2:** A time-lapse movie of migrating GFP^+^ cells within the dorsal region of a ventral/GFP::dorsal organoid fusion. This movie is an example of a cell exhibiting many changes of direction involving the dynamic extension and retracing of several processes. As the cell body remains static, branches are extended in multiple directions, and then each of the main branches extends additional higher order branches. Finally, a branch is extended in a particular direction followed by the retraction of the other main branch. The cell body is then moved in the direction of the extending branch. The cycle is repeated as the cell decides which direction to migrate.

**Supplemental Movie 3:** A time-lapse movie of migrating GFP^+^ cells within the dorsal region of a ventral/GFP::dorsal organoid fusion. This movie shows multiple migrating cells. 1) Initially a cell in the middle of the field of view is migrating upward. The upward process is retracted as a new leading process is extended downward and becomes branched. The cell migration direction is then changed downward. The bifurcated leading process is dynamic such that one process is extended as the other process is retraced. The cell body then moved toward the extended leading process. Prior to nucleokinesis, a swelling is observed moving from the cell body into the proximal portion of the leading process. Then the cell body is moved in parallel to the swelling, and finally the cell body moves into the swelling. 2) A second cell migrates from the left field of view toward the right, changes direction back toward the left, and then again changes direction toward the right, and finally changes once again back toward the left. With each change of direction, the trailing process becomes the leading process. The new leading process is extended toward the direction of travel as the trailing process is retracted.

**Supplemental Movie 4:** A time-lapse movie of migrating GFP^+^ cells within the dorsal region of a ventral/GFP::dorsal organoid fusion. This movie shows multiple cells migrating in different directions with extending and retracting processes. Beginning around 45 hrs, one cell migrates throughout the entire field of view beginning in the top left corner and traveling toward the bottom right corner. The cell travels rapidly in a constant direction, but at around 70 hours the progress is slowed as the leading process branches.

**Supplemental Movie 5:** A time-lapse movie of migrating GFP^+^ cells within the dorsal region of a ventral::dorsal organoid fusion. This movie shows multiple cells migrating. Around 6 hours a cell can be seen migrating into view from the bottom left corner of the field of view. This cell initially migrates left to right with a branched leading process. At multiple times, 3 branches are observed. As the cell progresses forward, branches are extended in the direction of travel, while other branches are retracted. Around 23 hours the cell changes direction abruptly from moving right to moving left. This involves an extension of a new process toward the left, while the previous leading process oriented to the right is retracted. The cell proceeds to the left, but around 39 hours, the leading process begins turning toward the right. The leading process makes a 180-degree turn, and then extends. The cell body then follows the leading process as the cell migrates rapidly from top to bottom and eventually proceeds out of view in the z-direction.

**Supplemental Movie 6:** A time-lapse movie of ventral-derived GFP^+^ neurites growing within the dorsal region of a ventral/GFP::dorsal organoid fusion. The neurites appear to be axons with an enlarged tuft at the end the processes which resembles that of a growth cone. The processes are highly dynamic, and exhibit extension in single directions, but also abrupt changes in direction.

